# HAND1 knockdown disrupts trophoblast global gene expression

**DOI:** 10.1101/2022.11.01.514704

**Authors:** Robert Fresch, Jennifer Courtney, Heather Brockway, Rebecca L. Wilson, Helen Jones

## Abstract

Congenital heart disease (CHD) affects nearly 1% of births annually, and CHD pregnancies carry increased risk of developing pathologies of abnormal placentation. We previously reported significant developmental impacts of disrupting *Hand1*, a gene associated with CHD, expression in placenta trophoblast and endothelial cells in multiple mouse models. In this study, we aimed to build upon this knowledge and characterize the mechanistic impacts of disrupting *HAND1* on human placenta trophoblast and vascular endothelial cell gene expression. *HAND1* gene expression was silenced in BeWo cells, a choriocarcinoma model of human cytotrophoblasts, (n=3-9 passages) and isolated human placental microvascular endothelial cells (HPMVEC; n=3 passages), with *HAND1* siRNA for 96 h. Cells were harvested, mRNA isolated and RNA sequencing performed using the Illumina NextSeq 550 platform. Normalization and differential gene expression analyses were conducted using general linear modeling in edgeR packages. Statistical significance was determined using a log2 fold change of >1.0 or <−1.0 and unadjusted p-value ≤0.05. Panther DB was used for overrepresentation analysis, and String DB for protein association network analysis. There was downregulation of 664 genes, and upregulation of 59 genes in BeWo cells with direct *HAND1* knockdown. Overrepresentation analysis identified disruption to pathways including cell differentiation, localization, and cell projection organization. In contrast, only 7 genes were changed with direct *HAND1* knockdown in HPMVECs. Disruption to *HAND1* expression significantly alters gene expression profile in trophoblast but not endothelial cells. This data provides further evidence that future studies on genetic perturbations in CHDs should consider the extra-embryonic tissue in addition to the fetal heart.

## Introduction

Congenital heart disease (CHD) affects nearly 1% of births annually, and often requires surgical intervention for repair and correction [1]. Despite advances in care, CHD is still a leading cause of infant mortality from pregnancies complicated by birth defects accounting for approximately 4.2% of all neonatal deaths, and is associated with healthcare costs of approximately $6.1 billion dollars annually [2]. About 25% of babies with a CHD will have a severe CHD [3], and the prevalence of CHD, particularly mild disease, is increasing. CHD pregnancies also carry increased risk of developing pathologies of abnormal placentation including fetal growth restriction (FGR), preeclampsia, preterm birth, and stillbirth [4]. These adverse pregnancy outcomes greatly impact cardiac care, specifically with regards to surgical morbidity and mortality, as well as impacting childhood development and survival [5].

Currently, we lack comprehensive understanding of the embryonic and fetal relationship between development of the placenta and the heart. Using a systematic computational approach, we have shown numerous commonly expressed genes between first trimester human heart and placenta cells, which if disrupted may concurrently contribute to the developmental perturbations resulting in CHD (Wilson 2022). Our lab has previously reported disrupted vascular development, and morphologic abnormalities and placental insufficiency in placentas from human pregnancies with CHD [5,6]. The placenta is essential to fetal growth and development and placental disfunction impacts perinatal outcomes [7]. This is because the placenta plays an essential role in regulating the transport of nutrients and oxygen from the mother to the fetus, and mediates maternal-fetal communication. In utero, initial heart and placenta development occurs in parallel during the first three weeks of gestation. We and others have shown common molecular pathways in placental and heart development [8,9]. However, in-depth knowledge of the regulation of these common molecular pathways, particularly in relation to vasculogenesis and angiogenesis, is lacking. Additionally, studies using mice models to better understand heart development have also been shown to exhibit abnormal placental development, although the latter is very rarely investigated [10].

*HAND1* is a transcription factor related to the basic helix-loop-helix (bHLH) with essential roles in embryonic placenta and heart development [11], and is expressed in first trimester human placental trophoblast [12,13] and first trimester cardiac cells [14]. Hand1-null mice are embryonic lethal by E8.5 due to defects in the extraembryonic tissues [11,15–17]. The Nkx2.5^Cre^/Hand1^A126fs/+^ mouse model is characterized by embryonic lethality at gestational day 15.5, as well as outflow tract abnormalities, thin myocardium and ventricular septal defects in the fetal heart [11]. More recently, we have shown that placentas of the Nkx2.5^Cre^/Hand1^A126fs/+^ mouse, in which *Hand1* is mutated in trophoblast progenitor cells, fail to develop the appropriate cell layers in the labyrinth (nutrient exchange area), resulting in fetal demise [8]. However, using the Cdh5^cre^/Hand1^A126fs/+^ mouse model, which specifically mutates *Hand1* in endothelial cells of the placenta and heart, the placentas are only affected in later-gestation with reduced placental vascular branching, but little effect on fetal heart development [8]. Signaling between trophoblast cells and villous endothelium is necessary for placental development and function, however there is a paucity of data looking at how signaling occurs at a molecular level during development. In this study, we aimed to build upon our discoveries in the mouse models and characterize the impact of disrupting *HAND1* expression on molecular signaling in human placenta trophoblast and placental villous endothelial cells.

## Materials and Methods

### BeWo Cell Culture

BeWo choriocarcinoma cells, which have physiological characteristics of the villous trophoblast [18,19], were maintained at 37°C, 5% CO_2_ in Ham’s F-12 medium (Sigma, St. Louis, MO) with 1% penicillin-streptomycin (Gibco, Waltham, MA), and 10% fetal bovine serum. Cells were sub-cultured every 3-4 days based on confluence estimates of 70-90%. Experiments were conducted on cells at passages 4-10.

### Human Placental Microvascular Endothelial Cell Culture

Human Placenta Microvascular Endothelial Cells (HPMVECs) were cultured in T75 flasks pre-treated with attachment factor (Cell Applications Inc.) at 37°C, 5% CO_2_ in EGM-2 media (Lonza, Allendale, NJ). Cells were sub-cultured every 3-4 days based on confluence estimates of 70-90% with media exchanges occurring every 2 days. Experiments were conducted on cells between passages 4-10.

### Direct *HAND1* knockdown in BeWo and HPMVECs

BeWo cells or HPMVECs were plated (2.5 × 10^5^ cells/well) onto Millicell hanging cell culture inserts (Millipore, Bedford, MA) in 12 chamber culture trays with respective culture media in both the well insert chamber. For HPMVECs, the inserts were pre-coated with attachment factor. After 24 h, the well culture media was removed, cells washed with PBS, and replaced with treatment media: minimum essential media (MEM; Sigma, St. Louis, MO) containing 1% L-glutamine (Gibco, Waltham, MA) and 1% penicillin-streptomycin. To knockdown *HAND1* cells (n = 9 passages for BeWo cells and n = 3 passages for HPMVECs) were treated with 3 µL Lipofectamine + 4 µL 10 µM *HAND1* siRNA for 96 h as laboratory standard [20]. Treatment of cells with 3 µL Lipofectamine + 3 µL 10 µM Allstars negative siRNA was used as a negative control. After 6 hours, 10% FBS was added to ensure cell survival without starvation effects of MEM. At 96 h, cells were harvested for RNA isolation following treatment.

### Isolation of RNA and confirmation of *HAND1* knockdown via QPCR

Cells were lysed using RLT Buffer from Qiagen (Valencia, CA) following manufacturer’s instructions. Total RNA was isolated using the RNeasy Mini Kit, QIAshredder, and on-column DNA digest (Qiagen) following the protocol provided by the manufacturer. For QPCR analysis, total RNA was quantified using a Nanodrop Spectrophotometer. 1 µg of RNA was then converted to cDNA utilizing the Applied Biosystems High Capacity cDNA kit following manufacturer’s protocol. QPCR was performed in duplicate reactions containing PowerUp SYBR Green (Applied Biosystems) as per manufacturer’s instructions and with primers (Supplemental Table 1) on the StepOne-Plus Real-Time PCR System (Applied Biosystems). Relative mRNA expression was calculated using the comparative CT method [21] with the StepOne Software v2.3 (Applied Biosystems) normalizing genes to *ACTB*

**Table 1:** Biological processes impacted by direct knockdown of the *HAND1* gene in BeWo cells

|  | Fold Enrichment | Adjusted P value<sup>a</sup><br>Bonferroni |
| --- | --- | --- |
| **GO Biological Process: Developmental Processes (GO:0032502)** |  |  |
| **Reactome pathways** |  |  |
| MET activates PTK2 signaling (R-HSA-8874081) | 17.2 | 4.11E-02 |
| Organelle biogenesis and maintenance (R-HSA-1852241) | 4.6 | 1.74E-02 |
| Signal Transduction (R-HSA-162582) | 1.89 | 1.73E-02 |
| **GO Biological Process: Cell Projection and Organization (GO:0030030)** |  |  |
| **Panther Pathways** |  |  |
| Organelle biogenesis and maintenance (R-HSA-1852241) | 10.2 | 1.47E-06 |
| Signaling by Rho GTPases (R-HSA-194315) | 6.04 | 5.16E-03 |
| **GO Biological Process: Regulation of Localization (GO:0032879)** |  |  |
| **Panther Pathways** |  |  |
| Integrin signaling pathway (P00034) | 5.29 | 2.79E-02 |
| **Reactome Pathways** |  |  |
| MET activates PTK2 signaling (R-HSA-8874081) | 21.06 | 1.56E-02 |
| Cardiac conduction (R-HSA-5576891) | 9.09 | 5.99E-04 |
| Muscle contraction (R-HSA-397014) | 6.08 | 1.87E-02 |
| **GO Biological Process: Establishment of Localization (GO:0051234)** |  |  |
| **Reactome pathways** |  |  |
| ABC-family proteins mediated transport (R-HSA-382556) | 7.16 | 1.81E-02 |
| Membrane Trafficking (R-HSA-199991) | 3.63 | 1.75E-05 |
| **GO Biological Process: Regulation of Multicellular Organismal Process (GO:0051239)** |  |  |
| **Panther Pathways** |  |  |
| TGF-beta signaling pathway (P00052) | 7.98 | 2.24E-02 |
| Gonadotropin-releasing hormone receptor pathway (P06664) | 7.86 | 9.78E-07 |
| EGF receptor signaling pathway (P00018) | 6.71 | 1.76E-02 |
| **Reactome pathways** |  |  |
| Toll-like Receptor Cascades (R-HSA-168898) | 8.57 | 9.80E-04 |
| Cardiac conduction (R-HSA-5576891) | 8.44 | 4.22E-03 |
| Toll Like Receptor 4 (TLR4) Cascade (R-HSA-166016) | 8.27 | 1.82E-02 |

<sup>a</sup> Statistical significance of over-representation was determined using Fisher's Exact Test with Bonferroni corrections for multiple corrections.

### Transcriptome generation and differential gene expression bioinformatic analyses

Total RNA was isolated from cells using same protocol as for QPCR. 50-100 μg of RNA from the various treated BeWo cells, and HPMVECs (n=3 passages submitted to the University of Cincinnati Genomics, Epigenomics and Sequencing Core for RNA quality assessment and sequencing. RNA quality control (QC)was conducted on an Agilent 2100 Bioanalyzer and all samples passed control checks with acceptable RNA integrity numbers (RIN). For each experimental data set, poly-A RNA libraries were generated using (NEBNext Ultra Directional RNA Library Prep kit, New England Biolabs, Ipswich, MA and the TruSeq SR Cluster kit v3 Illumina). Transcriptomes were generated on the Illumina NextSeq 550 platform with ∼25 million reads per sample with a single end read length between 85-101bp. Initial quality control for post-sequencing reads, read alignment, and read count generation were all performed in the public Galaxy Bioinformatic server [22] utilizing the following tools: FASTQC [23], trimmomatic [24], Bowtie2 [25], and featurecounts [26]. All samples were then aligned utilizing the hg38 genome build via Bowtie2, which allowed for more precise alignments of the numerous homologous genes expressed in these specific cell lines. For each different experiment, gene count matrices were generated using featurecounts and utilized for differential gene expression analyses. Differential gene expression analyses were conducted using the Empirical analysis of digital gene expression in R (EdgeR) package [27]. General linear modeling using the following pairwise comparisons were performed between, untreated controls, Allstars negative control treated, direct *HAND1* knockdown treated BeWo and HPMVEC cells that were treated directly. Multiple corrections testing yielded no statistical differences in the pairwise comparisons. Therefore, we used the raw p-values to determine genes to be used in overrepresentation analysis to identify pathways and processes rather than individual genes. RNA sequencing data have been deposited to NCBI GEO under the accession GSE209620.

### Overrepresentation analysis

#### Panther DB Evaluation

Lists of significantly differentially expressed genes identified between Allstar negative control and *HAND1* siRNA treated BeWos and HPMVECs were analyzed by PantherDB (Panther15.0) to determine over-representation and identify pathways and processes involved in trophoblast-endothelial cell signaling. Gene names were submitted with statistical testing conducted using Fisher’s Exact test with multiple corrections testing via Bonferroni correction. We conducted analyses using Panther pathways, Reactome pathways, and GO Biological Processes against the entire genome for Homo sapiens.

#### *In silico* StringDB assessment of interaction networks

StringDB (version 11.0) was utilized to assess potential protein interactions affected by knockdown of *HAND1* in BeWo cells. Seven significantly differentially expressed genes with large fold-changes were individually entered into StringDB and then functional interactions classified into biological pathways. Parameters were set at Homo Sapiens, Experimental and Database sources, Full Network Search.

### Statistical Analysis

qPCR data were analyzed in Prism v8 (GraphPad) using either Kruskal-Wallis test with Dunn’s multiple comparison test or Mann-Whitney test. Data for qPCR is presented as the median ± interquartile range.

## Results

### *HAND1* siRNA treatment knocked-down *HAND1* expression in BeWo cells and HPMVECs

Compared to Allstar negative control treated cells, treatment with *HAND1* siRNA for 96 h reduced *HAND1* expression 61% in BeWo and 69% HPMVECs (Figure 1A & 1B, respectively).

**Figure 1.**
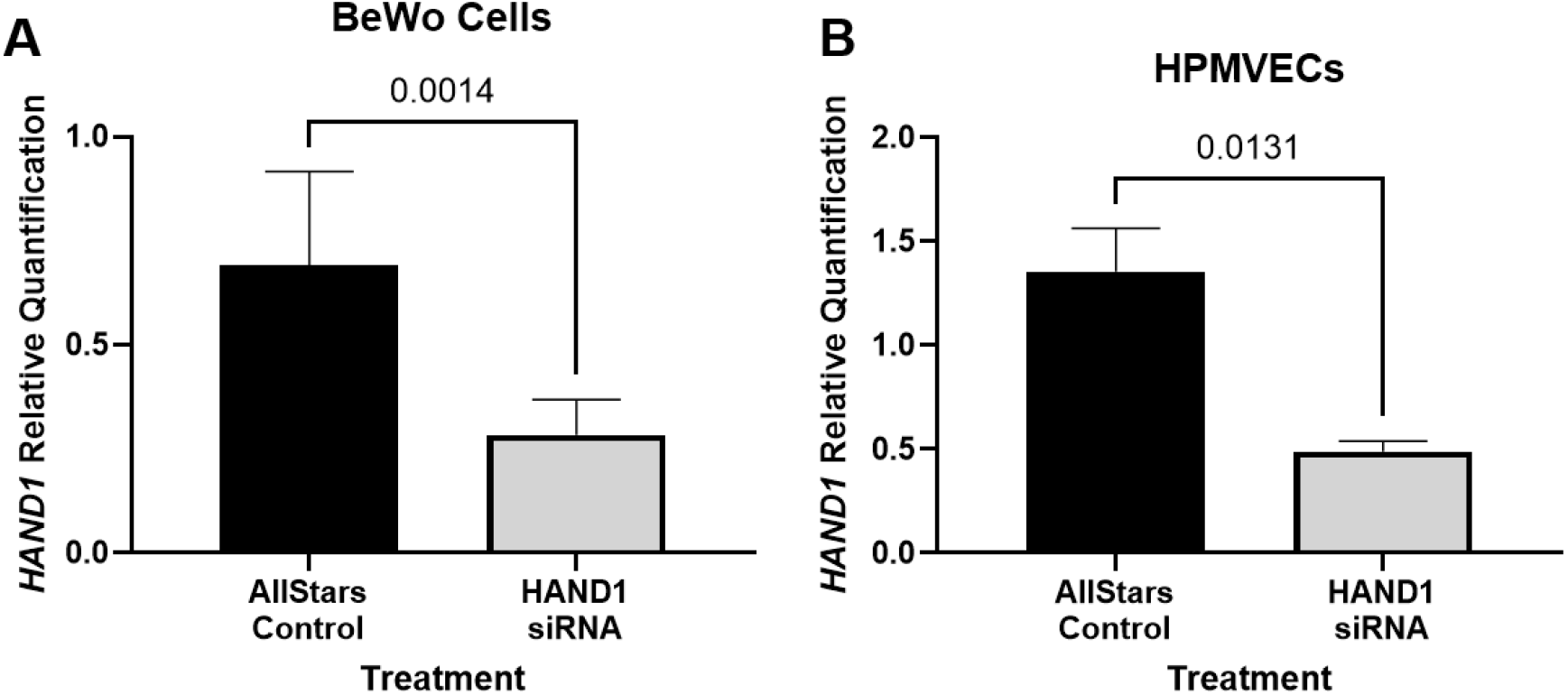
*HAND1* mRNA expression in BeWo cells and Human Placental Microvascular Endothelial Cells (HPMVECs) following siRNA treatment. **A**. In BeWo cells, *HAND1* mRNA expression was reduced by 61% compared to Allstar negative control treatment after 96 h. **D**. In HPMVECs, *HAND1* mRNA expression was reduced by 69% compared to Allstar negative control treatment after 96 h. Data are median ± interquartile range. Statistical significance was determined using Mann-Whitney test. n = 3 to 9 passages

### Direct *HAND1* siRNA treatment in BeWo cells resulted in changes to global gene expression

Compared to Allstar negative control, *HAND1* knockdown in BeWo cells resulted in downregulation of 664 genes, and upregulation of 59 genes (Supplemental Material). PantherDB was utilized to perform overrepresentation analysis against Panther pathways, Reactome pathways, and Gene Ontology (GO) Biological processes on the differentially expressed genes. No overrepresentation was seen compared to Panther and Reactome pathways, however many results were returned for GO Biological processes (Supplemental Table 2). Groups of genes identified as significantly over-represented in GO Biological processes were then re-entered into PantherDB and indicated potential disruption to pathways including cell development, cellular projection, regulation and establishment of localization, and regulation of multicellular function (Table 1). There were several biological pathways over-represented, including GnRH releasing hormone pathways, cardiac conduction and signaling, TGF-beta signaling, and signaling of RHO GTPases. In addition, there was significant enrichment in MET activating PTK2 signaling, and pathways related to signal transduction.

**Table 2:** Differentially expressed genes with large fold-change differences in BeWo cells in which HAND1 was knocked down

|  | Gene (Ensembl gene ID) | Log2 FC | Raw P-value<sup>a</sup> |
| --- | --- | --- | --- |
| **Upregulated** |  |  |  |
|  | *CALML5* (ENSG00000178372) | 4.6384 | 0.0012 |
|  | *NUBP1* (ENSG00000103274) | 4.2640 | 0.0124 |
|  | *TFAP2E* (ENSG00000116819) | 3.5473 | 0.0041 |
|  | *WNT8A* (ENSG00000061492) | 2.9174 | 0.0065 |
| **Downregulated** |  |  |  |
|  | *FAM49B* (ENSG00000153310) | -5.8251 | 0.0001 |
|  | *CTTNBP2* (ENSG00000077063) | -5.6647 | 0.0007 |
|  | *NFS1* (ENSG00000244005) | -5.4039 | 0.0001 |

<sup>a</sup> Statistical testing applied within edgeR general linear models. FC = fold-change

StringDB network analysis of seven genes with large fold change differences in expression following *HAND1* knockdown in BeWo cells were found to be involved in biological pathways with known importance in growth and development (Table 2). Upregulated genes were *CALML5, NUBP1, TFAP2E*, and *WNT8A*. Downregulated genes; *FAM49B, CTTNBP2*, and *NFS1*. Reactome pathways examined identified common relationships between the genes including Beta-catenin phosphorylation, TGF-beta signaling, and Pl3K-Akt signaling. Other notable biological pathways included cardiac conduction and calcium channel signaling, GnRH and Estrogen dependent gene expression, eNOS activation and regulation, and iron-sulfur and sulfur metabolism pathways and RHO GTPases (Figure 2).

**Figure 2.**
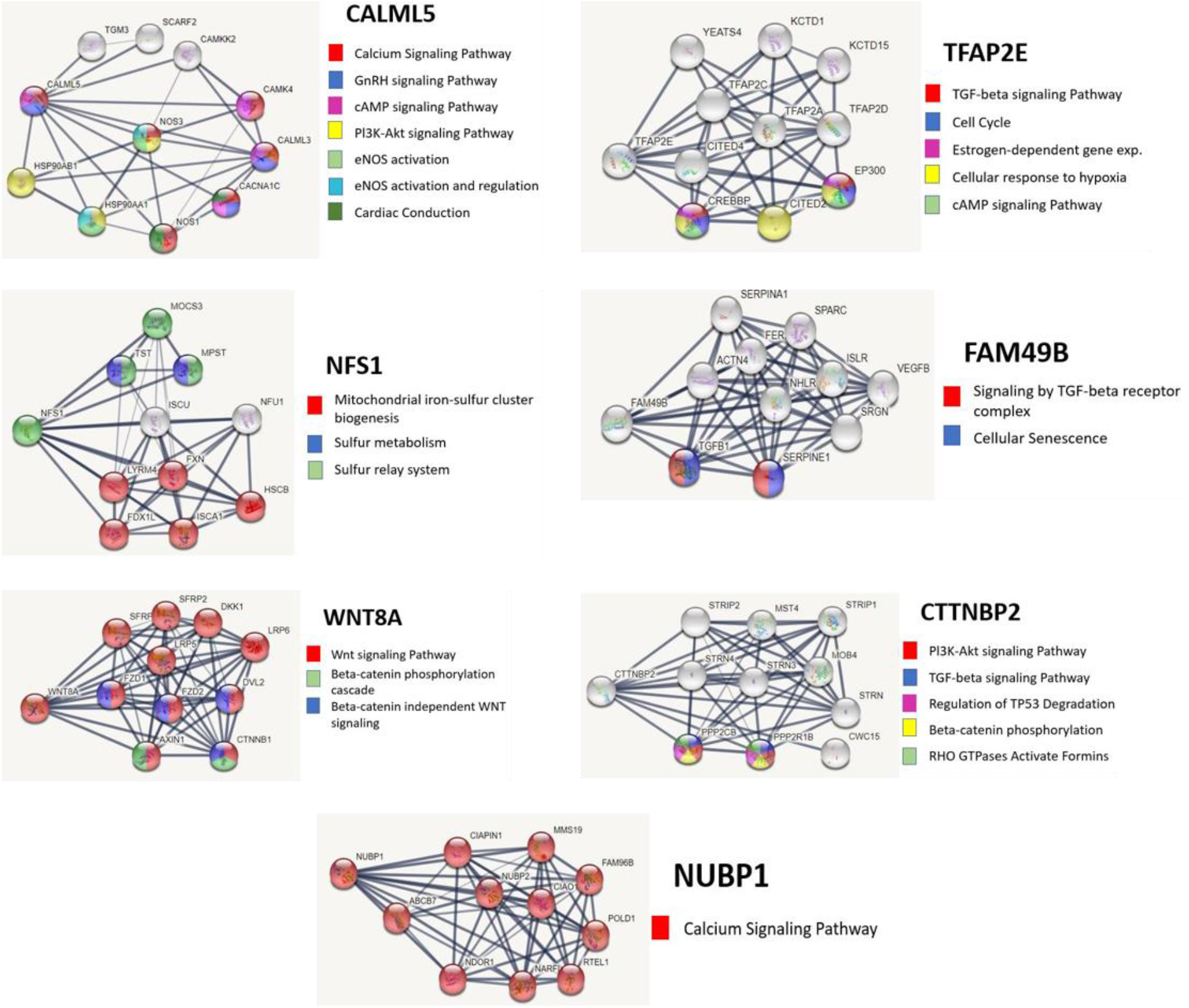
Network analyses of genes disrupted by HAND1 knockdown in BeWo. Expression of *CALML5* (**A**), *NUBP1* (**B**), *TFAP2E* (**C**), and *WNT8A* (**D**) was significantly upregulated in BeWo cells treated with *HAND1* siRNA when compared to Allstar negative control. Expression of *FAM49B* (**E**), *CTTNBP2* (**F**), and *NFS1* (**G**) was significantly downregulated in BeWo cells treated with *HAND1* siRNA when compared to Allstar negative control. Grey lines represent the network edges with thickness representing the confidence of the data support (thicker lines = higher confidence data). Color coded legends show genes in Reactome pathways that may be impacted by *HAND1* knockdown.

### Direct treatment of HPMVECs with *HAND1* siRNA minimally disrupted global gene expression

Direct *HAND1* knockdown in HPMVECs resulted in minimal disruption to global gene expression with differential expression in just seven genes (Table 3). QPCR validation confirmed two genes of interest, *GADD45g* and *NPPB* as reduced and increased, respectively in *HAND1* siRNA treated HPMVECs when compared to Allstar negative control (Figure 3).

**Table 3.**
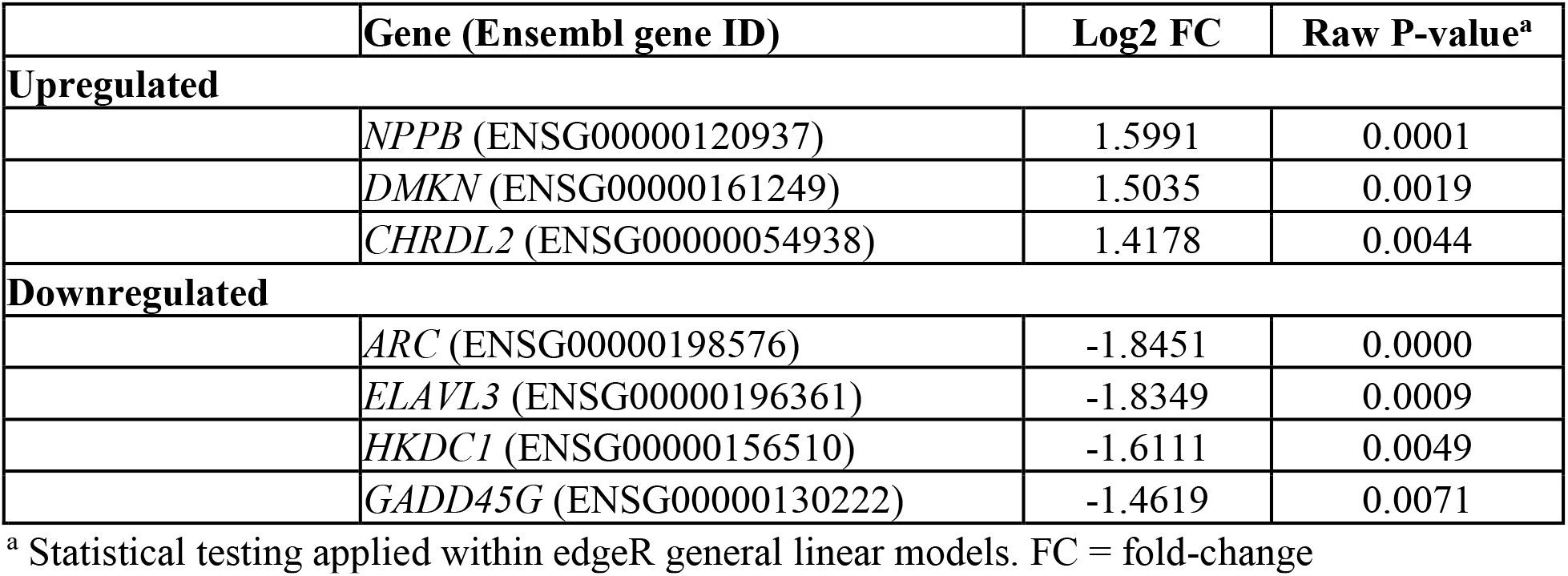
Significantly differentially expressed genes in Human Placenta Microvascular Endothelial cells in which HAND1 was knocked down

**Figure 3.**
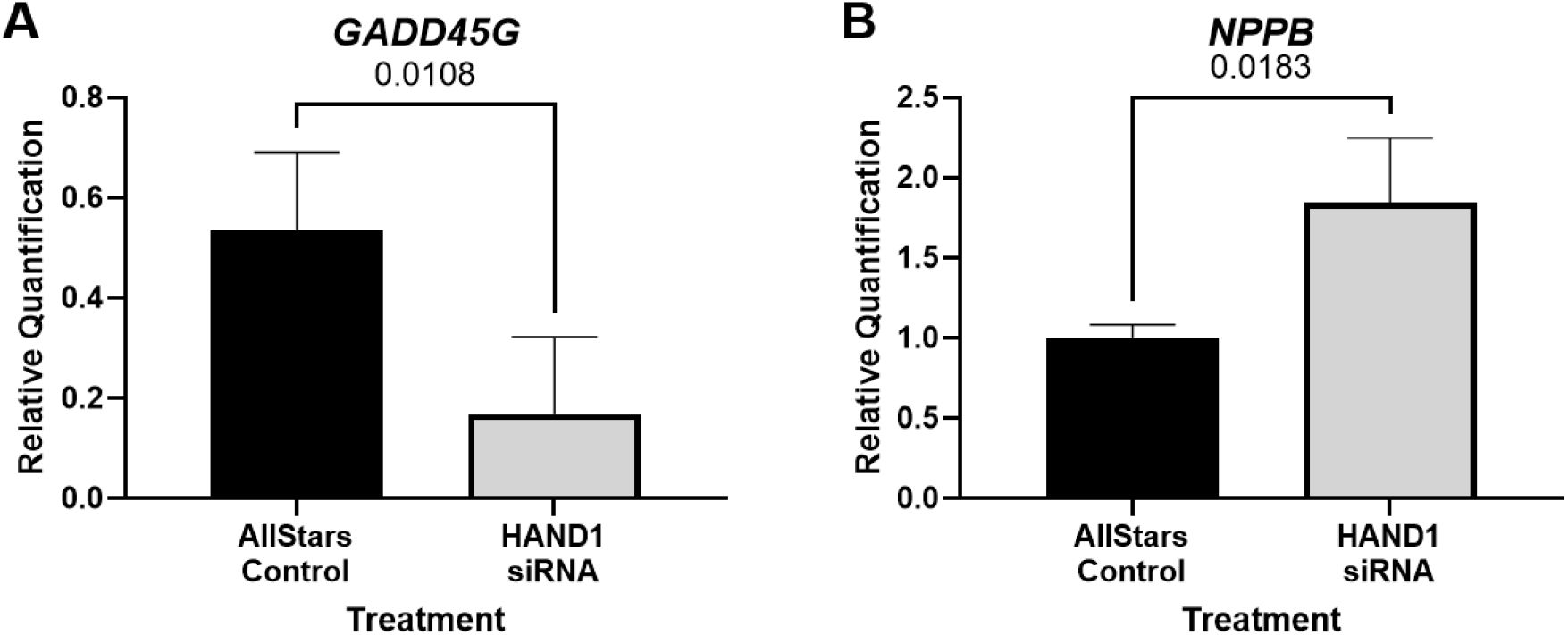
qPCR validation of two genes shown to be differentially expressed using RNA sequencing. **A**. mRNA expression of *GADD45G* was shown to be reduced in *HAND1* siRNA treated Human Placenta Microvascular Endothelial cells (HPMVECs) when compared to Allstar negative control treated cells. **B**. Expression of *NPPB* was increased in in *HAND1* siRNA treated HPMVECs when compared to Allstar negative control treated cells. Data are median ± interquartile range. Statistical significance was determined using Mann-Whitney test. n = 3 passages

## Discussion

CHDs are often associated with pregnancy complications such as fetal growth restriction and preeclampsia, these conditions negatively impact clinical outcomes, increase the risk of neonatal morbidity and mortality, and are likely a consequence of abnormal placental development and function [28–32]. We have previously shown in mice, that targeted loss of *Hand1* in chorionic and labyrinthine progenitor trophoblasts led to abnormal formation of the placental labyrinth, and ultimately embryonic lethality [8]. Histological analysis of the placenta indicated that loss of *Hand1* in labyrinthine progenitor trophoblasts early in pregnancy significantly impacted the ability for the placenta to form syncytial layers, and impacted development of the labyrinthine vasculature. In this study we aimed to gain further mechanistic, translational understanding of the effects of *HAND1* knockdown in models of human placenta trophoblast and villous endothelial cells. We demonstrated significant alterations to placental trophoblast gene expression following *HAND1* knockdown, and identified potential pathways which may be significantly impacted by loss of *HAND1* regulation. This study is the first to identify possible molecular signaling pathways that are impacted by disruption to *HAND1* in the human placenta.

Overall, there were 664 genes differentially expressed in BeWo cells due to *HAND1* knockdown. Overrepresentation analyses reveals several key GO Biological Processes including: cell development, establishment of localization, and regulation of multicellular function, as well as biological pathways including Pl3K-Akt signaling, signaling Rho GTPases, and TLR cascades. Trophoblast differentiation during the first trimester of pregnancy involves trophoblast proliferation, invasion and extracellular matrix (ECM) remodeling. PI3K/Akt signaling reduction or inhibition plays an important role in trophoblast proliferation, migration, and survival. Disruption to PI3K/Akt signaling in early embryonic development is associated with growth restriction, preterm birth, and embryonic lethality [33], highlighting the importance of this signaling pathway to placental development and function. Additionally, inhibition of PI3K increases soluble fms-like tyrosine kinase 1 (sFlt1), a common biomarker of pre-eclampsia [34]. PI3K/Akt signaling has been closely linked to signaling Rho-GTPases which are known to play a role in trophoblast migration [35,36].

TLR cascades form the major family of pattern recognition receptors that are involved in innate immunity. The maternal-fetal interface immunologically is unique in that it must promote tolerance of the fetus while maintaining protection to the mother. Trophoblasts play an important role in modulating the maternal immune response throughout pregnancy, including through TLR signaling [37]. Additionally, TLR signaling has been shown to potentially modulate angiogenesis as culture of trophoblasts with TLR2 ligand HKML have been shown to promote the expression of pro-angiogenic Placenta Growth Factor [38]. Overall, poor migration of trophoblasts, and communication with resident immune cells, can impact invasion and establishment of a fully functional maternal-fetal interface.

Expression of *CALML5* and *NUBP1* was upregulated in BeWo cells following *HAND1* knockdown. Both genes are involved with Calcium channel signaling, GnRH signaling, cAMP signaling, eNOS activation and Pl3K-Akt signaling pathways. These are important biological pathways that impact trophoblast invasion, differentiation, development, resource control and growth of the placenta and fetus [39,40], and increased gene expression of *CALML5* and *NUBP1* may be a compensatory response to disruption of other signaling pathways. On the other hand, expression of *CTTNBP2* and *NSF1* was downregulated in BeWo cells in which *HAND1* was knocked down. *CTTNBP2* has been shown to have a direct relationship with the WNT signaling pathway [41], and downregulation in WNT signaling in the placenta has been associated with pathological pregnancies [42]. Similarly, *NFS1* is a gene that has an essential role in iron-sulfur cluster processing making it important for electron transport, enzyme catalysis, and regulation of gene expression as well is iron homeostasis [43]. Fetal growth is very dependent on energy metabolism in the placenta as it drives exchange of nutrients and plays a crucial role in DNA synthesis. Overall, our data indicates potential disruption to these pathways with *HAND1* knockdown and provides further understanding of how a genetic perturbation in this gene may lead to growth issues, developmental defects, and lethality/miscarriage in the context of human pregnancies with CHDs.

We sought to analyze the effect of *HAND1* knockdown in cells within the villous environment. Interestingly, direct *HAND1* knockdown in villous endothelial cells resulted in minimal impact to gene expression. This result however is in agreement with our mouse model studies suggest that disrupting *Hand1* expression directly in labyrinthine endothelial cells impacted vascular remodeling only in late pregnancy and non-branching angiogenesis mechanisms[8] not as individual endothelial cells or vasculogenesis. HPMVECs are cultured as a single monolayer. Therefore, it would be interesting for future studies, beyond the scope of the current study, to assess angiogenesis and remodeling mechanisms in a 3D vascularized model when HAND1/2 is knocked down or cultured in ‘conditioned’ media from BeWo cells treated with *HAND1/2* siRNA. Cell-cell communication/signaling within the placenta villi in the human is believed to be important in establishment of the villous structure and exchange region but given our current data, the involvement of other cell types such as stromal fibroblasts in the communication process requires future investigation.

We and others have consistently shown that *HAND1* is important to both fetal heart and placenta development [8,9], with the present study providing further mechanistic understanding of how *HAND1* may influence the development of the placenta in the human. Given our data shows greater disruption to global gene expression in placenta trophoblasts then endothelial cells with *HAND1* knockdown, this further highlights the importance of future research to consider analyzing the extra-embryonic tissue, as well as the heart, in the context of CHD.

